# Hi-C Calibration by Chemically Induced Chromosomal Interactions

**DOI:** 10.1101/2024.12.09.627644

**Authors:** Yi Li, Fan Zou, Lu Bai

## Abstract

The genome-wide chromosome conformation capture method, Hi-C, has greatly advanced our understanding of genome organization. However, its quantitative properties, including sensitivity, bias, and linearity, remain challenging to assess. Measuring these properties *in vivo* is difficult due to the heterogenous and dynamic nature of chromosomal interactions. Here, using Chemically Induced Chromosomal Interaction (CICI) method, we create stable intra- and inter-chromosomal interactions in G1-phase budding yeast across a broad range of contact frequencies. Hi-C analysis of these engineered cell populations demonstrates that static intra-chromosomal loops do not generate Topologically Associated Domains (TADs) and only promote 3D proximity within ∼50kb flanking regions. At moderate sequencing depth, Hi-C is sensitive enough to detect interactions occurring in 5-10% of cells. It also shows no inherent bias toward intra-versus inter-chromosomal interactions. Furthermore, we observe a linear relationship between Hi-C signal intensity and contact frequency. These findings illuminate the intrinsic properties of the Hi-C assay and provide a robust framework for its calibration.

## Introduction

Chromosome conformation capture-based methods, such as Hi-C, evaluate spatial proximity between distant chromosomal regions by measuring ligation frequencies (1, 2). While Hi-C signals are generally interpreted as “contact frequencies,” their quantitative interpretation remains challenging (3). Specifically, Hi-C-derived contact frequencies cannot be directly converted into the fraction of cells that engages in these interactions, and changes in interaction levels may not result in proportional changes in Hi-C signals. Furthermore, there may be biases in Hi-C signals among loci pairs. For example, it was reported that inter-chromosomal interactions are underrepresented in Hi-C data compared to intra-chromosomal ones, even though they occur at similar frequencies (4).

In principle, Hi-C can be calibrated using imaging techniques such as DNA Fluorescence In Situ Hybridization (FISH), i.e. by visualizing loci pairs and comparing their co-localization probabilities with the corresponding Hi-C signals (5). However, due to limited optical resolution, imaging-based quantification of co-localization relies on thresholding, making it difficult to accurately measure the fraction of cells where the two loci interact. To address this issue, we took advantage of Chemically Induced Chromosomal Interaction (CICI), a synthetic biology method we previously developed in budding yeast (6). By expressing LacI-FKBP12 and TetR-FRB fusion proteins that are targeted to integrated LacO and TetO arrays, stable interactions can be induced between the two arrays upon rapamycin addition (**Figure 1A**). This system allows for accurate determination of the fraction of cells in which the loci pair are in contact, providing a means to calibrate Hi-C signals with absolute contact frequencies.

**Figure 1.**
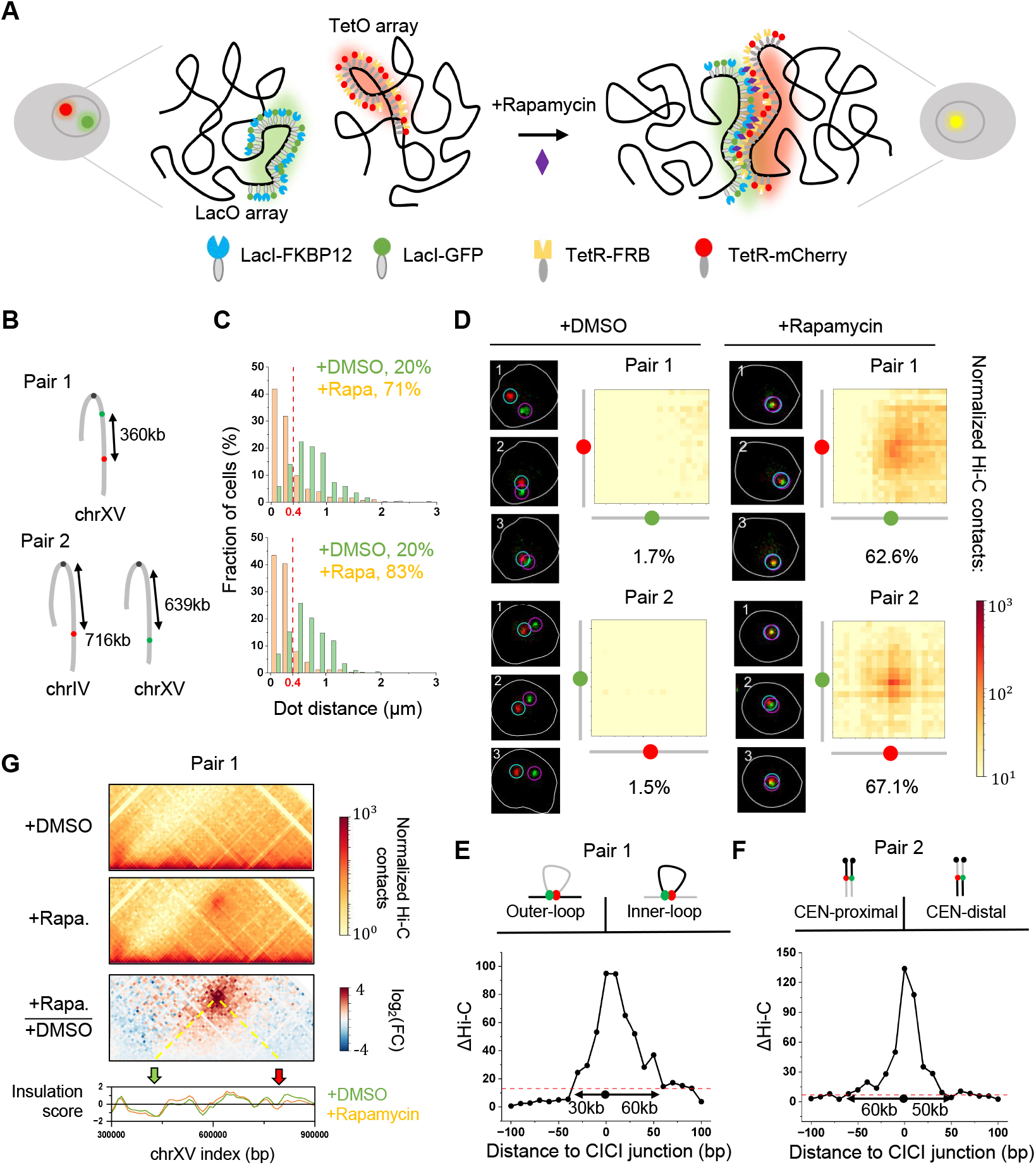
Imaging and Hi-C analysis of CICI. **A)** Schematic representation of the CICI system. LacO and TetO arrays are inserted into two chromosomal loci, and four fusion proteins are expressed to induce and visualize chromosomal interactions upon rapamycin addition. **B)** Configuration of LacO and TetO arrays for CICI pair 1 & 2. **C)** Histograms of dot distances between two loci pairs ± rapamycin. Threshold for co-localization (verticle line): 0.4 μm. Number of cells analyzed: pair 1 (+rapamycin: n = 257, -rapamycin: n = 286); pair 2 (+rapamycin: n = 346, - rapamycin: n = 202). **D)** Representative imaging data and Hi-C contact maps near the CICI junctions (± 200kb) ± rapamycin for both loci pairs. Images show three consecutive frames, with loci pairs considered to be in contact if dots are continuously co-localized. **E & F)** Differences in Hi-C signals ± rapamycin on both sides of CICI junctions. Difference above three standard deviation from the mean (marked by the red dotted lines) are considered as significant. Data in panels (C–G) are derived from two independent biological replicates. **G)** Static loops do not result in the formation of TADs. Normalized Hi-C contacts, log_2_ fold-change, and insulation scores ± rapamycin are shown.

In this study, we used CICI to generate cell populations with variable degrees of interaction between selected loci pairs. We then applied Hi-C to these cells to evaluate the sensitivity, bias, and linearity of the method. Our results show that, with 18M contact reads, Hi-C is sensitive enough to detect chromosomal interactions occurring in as few as ∼8% of cells. Moreover, for the loci pairs we examined, Hi-C does not display significant bias toward intra- vs inter-chromosomal interactions. We also find that static intra-chromosomal loops do not lead to the formation of TADs. Finally, our results demonstrate a linear relationship between Hi-C signal strength and the fraction of cells engaged in chromosomal interactions. Together, these findings provide insights into the nature of the Hi-C signals and their relation to genomic interactions.

## Results and Discussion

The CICI method relies on the expression of four fusion proteins, LacI-FKBP12, TetR-FRB, LacI-GFP, and TetR-mCherry, which bind to inserted LacO and TetO arrays in a rapamycin-resistant yeast strain. Upon rapamycin addition, dimerization of FKBP12 and FRB enables the two arrays to associate when in close proximity (**Figure 1A**). LacI-GFP and TetR-mCherry label the two arrays as distinct fluorescent foci, allowing direct visualization of CICI formation (6). In this study, we applied CICI to one intra-chromosomal loci pair (#1) and one inter-chromosomal loci pair (#2) in G1-arrested cells (**Figure 1B**). CICI was successfully induced for both loci pairs, as indicated by increased co-localization between the fluorescent foci in the presence of rapamycin (**Figure 1C**). Without rapamycin, ∼20% of cells exhibited co-localization with distance between the two arrays below the threshold 0.4 μm. In the presence of rapamycin, this number increases to 71.2% and 82.7% for the two pairs, respectively. To verify that observed co-localization represents genuine chromosomal interactions, we performed time-lapse imaging and calculated the absolute contact frequency as the fraction of cells that display continuous co-localization over time (**Materials and Methods**). This frequency is 62.6% and 67.1% for loci pair 1 and 2 with rapamycin, compared to merely 1.7% and 1.5% without it (**Figure 1D**), indicating that CICI can induce stable chromosomal interactions in 60.9% and 65.6% of cells. Note that the time-lapse approach reduces the impact of thresholding on the co-localization estimates. For example, extending the threshold to 0.5 μm only causes mild increases in CICI probabilities to 65.4% and 70.1%, respectively.

Next, we performed Hi-C measurements in the same cell populations with and without rapamycin treatment. The formation of CICIs for both locus pairs in the presence of rapamycin is evident from elevated Hi-C signals near the CICI junctions (**Figure 1D**). Specifically, local Hi-C contacts increased by 12-fold for pair 1 and 13-fold for pair 2, highly correlated with the corresponding increase in CICI probabilities. These findings suggest that, at least for contacts medicated by CICI, Hi-C does not show preference in detecting intra-chromosomal interactions. CICI formation also causes perturbations to the local chromatin architecture. For pair 1, ∼60 kb inner-loop region and ∼30 kb outer-loop region near the junction show significantly elevated proximity following CICI formation (**Figure 1E**). For pair 2, we observed increased trans-chromosomal interactions spanning 50-60kb regions flanking both sides of the CICI junction (**Figure 1F**).

As chromatin loops are often associated with TADs (7), we asked whether the CICI-induced loop at pair 1 leads to TAD formation. The short-range effect observed in **Figure 1E** cannot account for the length scale of TADs in mammalian cells, which typically span hundreds of thousands to a few million bps (2). To explore this further, we applied a standard TAD detection algorithm (8) to calculate the insulation scores for regions near pair 1, which revealed no differences with or without rapamycin (**Figure 1G**). This finding is consistent with previous polymer simulations showing that TADs emerge from dynamic loop extrusion rather than static loops (9). More specifically, during asynchronized loop extrusion, loops form at various locations among individual cells, bringing different regions within TAD boundaries together. The elevated proximity within the entire TADs is thus a consequence of population-averaging, rather than the whole regions forming highly compacted clusters.

To evaluate the sensitivity and linearity of Hi-C signals, we prepared cell populations ± CICI induction for both loci pairs, crosslinked them with formaldehyde, combined them in different ratios, and performed Hi-C measurements (**Figure 2A**). This approach allowed us to create artificial cell populations with contact frequencies ranging from 1.5% to 67.1%. The Hi-C results demonstrate that, with our modest sequencing depth of 18M contact reads, interactions in 7.8% of cells can be detected (**Figure 2B**). Remarkably, Hi-C signals have a largely linear relationship with contact frequencies across the full range (**Figure 2C**). This relation is essentially the same for both pairs, further suggesting that Hi-C has comparable sensitivity for intra- vs inter-chromosomal interactions. Using the fitted linear function, we converted the Hi-C signals measured over the native genome into absolute contact frequencies and averaged among regions separated by varying distances (**Figure 2D**). Regions within 40 kb show contact frequencies reaching 100%, indicating that contacts in this range are consistently captured by Hi-C. 40 kb genomic distance corresponds to 250–350 nm in 3D space (10), agreeing well with the estimated Hi-C capture radius of 100–400 nm (11). For regions that are separated by over 400kb, the interactions on average only occur in less than 1% of cells.

**Figure 2.**
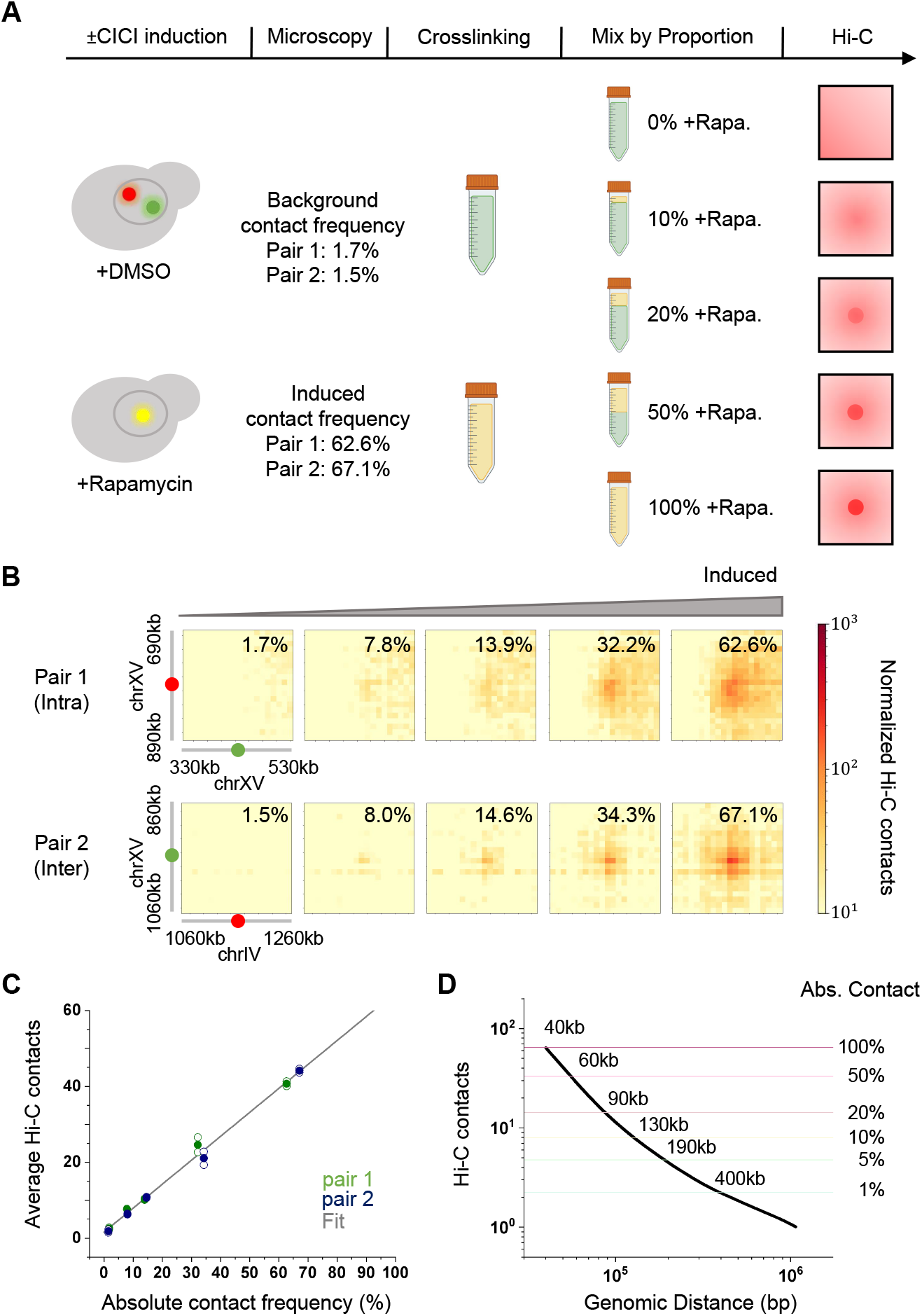
Hi-C signal scales linearly with CICI contact frequency. **A)** Experimental workflow. Populations of cells with variable contact frequencies between CICI pairs are generated by mixing fixed cells ± rapamycin at different ratios. **B)** Hi-C signals for each population in A. ± 200kb regions centered at the CICI junctions are shown. Data were merged from two biological replicates, and the absolute contact frequencies between loci pairs are indicated. **C)** Normalized Hi-C read counts as a function of contact frequencies between loci pairs. Hi-C signals are calculated by taking an average of ±10kb regions centered at the CICI junctions. Hollow and solid dots represent data from individual replicates and their mean, respectively. A linear fit is shown as a grey line. **D)** Hi-C contacts as a function of genomic distance based on the linear function in C. Averaged data from four - rapamycin measurements are used.

In comparison with the native genome, where the Hi-C signals and contact frequencies are hard to quantitatively interpret, the synthetic CICI system offers several advantages: 1) strong interactions can be induced between specific loci pairs in a large fraction of cells, 2) the stable nature of these interactions allows for reliable quantification of contact frequencies through time-lapse imaging, 3) interactions across different intra- and inter-chromosomal loci pairs can be created through the same mechanism, and 4) by mixing fixed cells with and without CICI at variable ratios, we can precisely modulate contact frequency while maintaining a consistent interaction mechanism. In this well-controlled system, Hi-C signals display the same linear relationship with contact frequency for both intra- vs inter-chromosomal loci pairs. Therefore, the reported non-linearity between contact frequency and Hi-C signals (12), as well as biases towards intra-chromosomal interactions (4), are not intrinsic limitations of the Hi-C methodology. Rather, the non-linearity and bias likely arise from the complexities in estimating “contacts” in native chromosomal contexts, where interactions are highly dynamic and vary in both distance and duration.

## Materials and Methods

### Plasmid and strain construction

All the strains used in this study were derived from a background strain carrying *tor1* and *fpr* mutations for rapamycin resistance. We constructed the CICI strain by integrating 1) plasmids containing LacO or TetO arrays into desired loci pairs, 2) a plasmid containing *LacI-FKBP12* and *TetR-FRB* driven by *REV1pr* into the *ADE2* locus, and 3) a plasmid containing *LacI-GFP* and *TetR-mCherry* driven by *REV1pr* into the *HIS3* locus. The tables below list all the plasmid and strains used here.

| Plasmid | Note | Note |
| --- | --- | --- |
| pSR11 | 192X TetO with a TRP1 marker | Rohner et al.(13) |
| pSR13 | 256X LacO with a LEU2 marker |  |
| pMY63 | REV1pr-LacI-GFP-REV1pr-TetR-mCherry with a HIS3 marker | Du et al.(6) |
| pMY48 | REV1pr-LacI-FKBP12-REV1pr-TetR-FRB with an ADE2 marker |  |
| pSR11-S1 | pSR11 plasmid with homologous sequence from Chr15::433394-433698 | This study |
| pSR11-S3 | pSR11 plasmid with homologous sequence from Chr15::793783-794105 |  |
| pSR11-S4 | pSR11 plasmid with homologous sequence from Chr15::965998-966332 |  |
| pSR11-S6 | pSR11 plasmid with homologous sequence from Chr4::1166520-1166835 |  |
| Strain | Genotype | Note |
| Pair 1<br>(yFZ102) | MAT<sub>a</sub>, *tor1-1*, *fpr::NAT*, *bar1*, *HIS3::REV1pr-tetR-mCherry-REV1pr-LacI-GFP*, *ADE2::REV1pr-LacI-FKBP12-REV1pr-TetR-FRB*, *Chr15::433546_LacO*, *Chr15::793958_TetO* | Intra pair |
| Pair 2<br>(yFZ110) | MAT<sub>a</sub>, *tor1-1*, *fpr::NAT*, *bar1*, *HIS3::REV1pr-tetR-mCherry-REV1pr-LacI-GFP*, *ADE2::REV1pr-LacI-FKBP12-REV1pr-TetR-FRB*, *Chr15::966175_LacO*, *Chr4::1166682_TetO* | Inter pair |

### CICI induction

Yeast strains were grown in 200 mL SCD-Met medium until OD660 reached 0.3. To arrest cells in the G1 phase, 80 μM alpha-factor (Zymo, Y1001) was added to the culture for a total of 2.5 h. During the final hour of this incubation, 1 ng/μL rapamycin (Sigma, 53123-88-9) was added to induce CICI. After 1 h of rapamycin treatment, cells were harvested for imaging analysis and Hi-C measurement.

### Imaging analysis

We performed time-lapse imaging with Leica fluorescent microscope (Leica DMI6000 with Hamamatsu ORCA-R2 C10600 camera), We used 7 z-stacks with 0.6 μm spacing for GFP and mCherry channels and 0.12∼0.15 s fluorescence excitation exposure was used for each stack. The time spent on each channel is about 18 s. Three consecutive time frames were taken with a time interval of 240 s. We used custom Matlab software developed previously to annotate cells, detect dot positions, and calculate dot distances (14). Absolute contact frequency for each loci pair is defined as the fraction of cells that show co-localized dots (<0.4μm) in three consecutive time frames.

### Hi-C experiments

Hi-C was adapted from a previously published protocol (15). Yeast were incubated in 200 mL SCD-Met medium until OD660 reached 0.3. Cells were fixed with 3% formaldehyde (Fisher, RSOF0010-250A) for 20 min at 25 °C and then quenched with 0.2 M glycine (Fisher, 194825) for 20 min at room temperature. Cells were collected by centrifugation and washed with SCD-Met. Populations of cells with different absolute contact frequencies between loci pairs were generated by mixing ± rapamycin populations by different proportions. Cell pellets were resuspended in 1 mL TBS (Tris-buffered saline), 1% Triton X-100 (Sigma, 9036-19-5) and 1X protease inhibitor cocktail (Thermo Fisher, 87786). Cell lysis was generated by adding 500 μL acid-washed glass beads (Sigma, G8772) and vortexing for 25 min at 4 °C. The chromatin was recovered through centrifugation, washed with 1 mL TBS, resuspended in 500 μL 10 mM Tris-HCl buffer and digested with DpnII (NEB, R0543L) overnight at 37 °C. Digested DNA fragments were filled in with biotin-labeled dATP by incubating with Klenow enzyme (NEB, M0212), biotin-14-dATP (Thermo Fisher, 19524016), dCTP, dTTP, and dGTP (Thermo Fisher, R0181) for 4 h at room temperature. The biotin-filled DNA fragments were ligated by T4 DNA ligase (NEB,M0202L) for 4 h at 16 °C. Crosslink was reversed by incubation with proteinase K (Thermo Fisher, EO0492) at 65 °C overnight. DNA was purified by phenol-chloroform extraction. Biotin-labeled, un-ligated fragment ends were removed by incubating with T4 DNA Polymerase (NEB, M0203), dATP and dGTP for 4 h at 20 °C. DNA was cleaned by DNA clean and concentrator-5 kit (Zymo, D4014) and sheared by Diagenode Biorupter Pico (EZ mode, 30 s on, 30 s off, 15 cycles). Biotin-labeled DNA was enriched by MyOne™ streptavidin C1 beads (Thermo Fisher, 65001). Hi-C libraries were prepared with NEBNext Ultra II DNA Library Prep Kit. Next generation sequencing was performed on NextSeq 2000, and 100 million of 50 bp paired-end reads were generated for each replicate. Hi-C data analysis was performed with HiC-pro (version 3.1.0) (16), cooler (version 0.10.0) (17), cooltools (version 0.7.0) (18) and HiCExplorer (version 3.7.2) (8).

## Data Availability

The sequencing data generated in this study have been deposited into the Gene Expression Omnibus (GEO) database under accession code GSE283767.

## Acknowledgments

We acknowledge all members in the Bai lab for insightful comments on the manuscript. This work is supported by the National Institutes of Health (T32 GM125592 to Y.L. and R35 GM139654 to L.B.) and the National Science Foundation (MCB-2016266 to L.B.).

